## Supplementary Information for "A global lipid map defines a network essential for Zika virus replication"

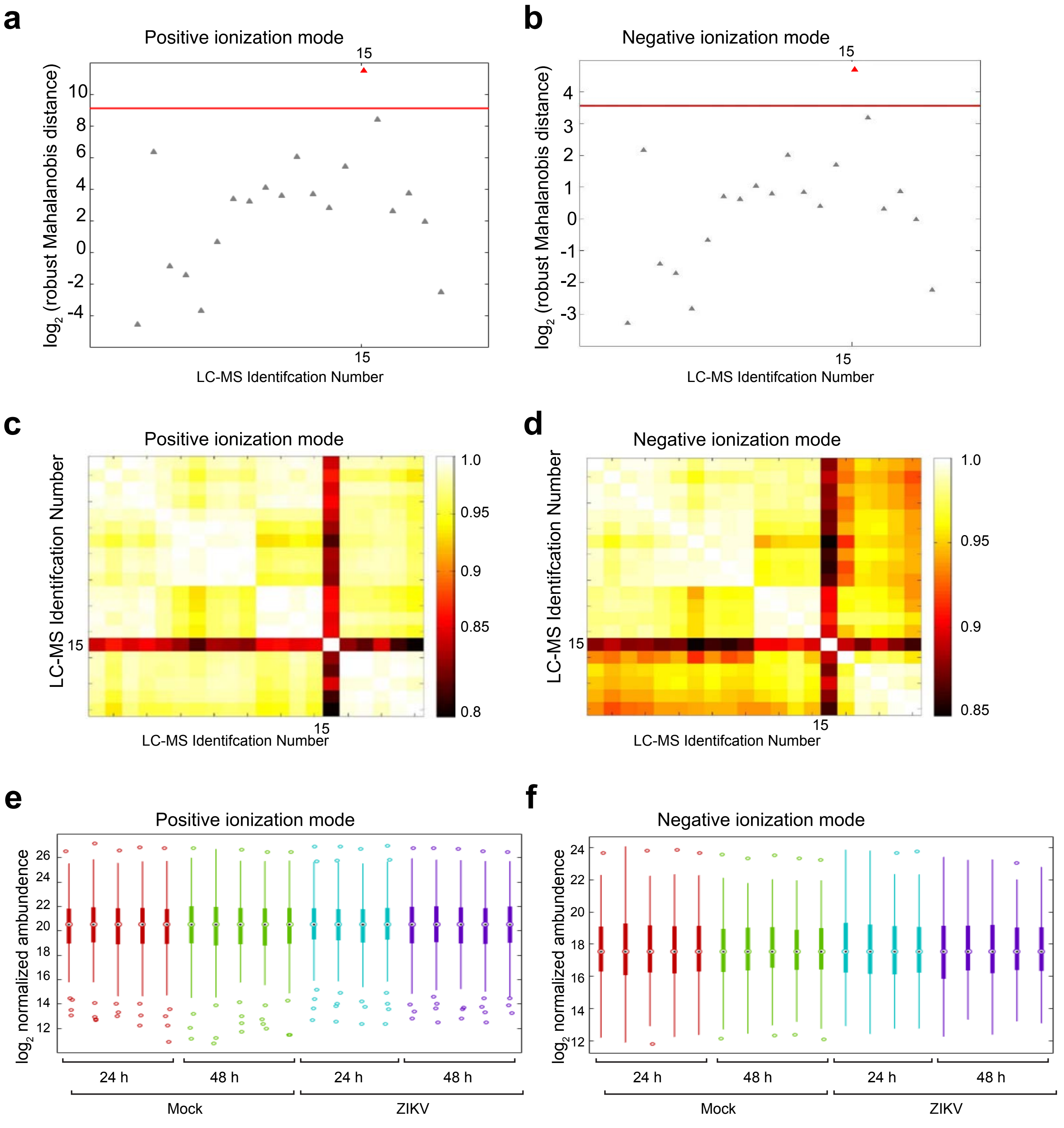


**Supplementary Fig. 1** Quality control and pre-processing of lipidomic data sets. **a**, **c**, **e** The robust Mahalanobis distance (RMD) analysis from instrument runs in positive ionization mode; **b**, **d**, **f** are from instrument runs in negative ionization mode. **a**, **b** RMD-PAV identifies LC-MS datasets that are extreme deviants from the remaining datasets (infected, 24 hpi; sample #15). **c**, **d** Heatmap of Pearson correlations confirms the presence of a single outlier sample (infected, 24 hpi; sample #15). We excluded sample #15 from further analysis. **e**, **f** Standard boxplot of the full mass spectrometry dataset after normalization. See also Supplementary Data 1.

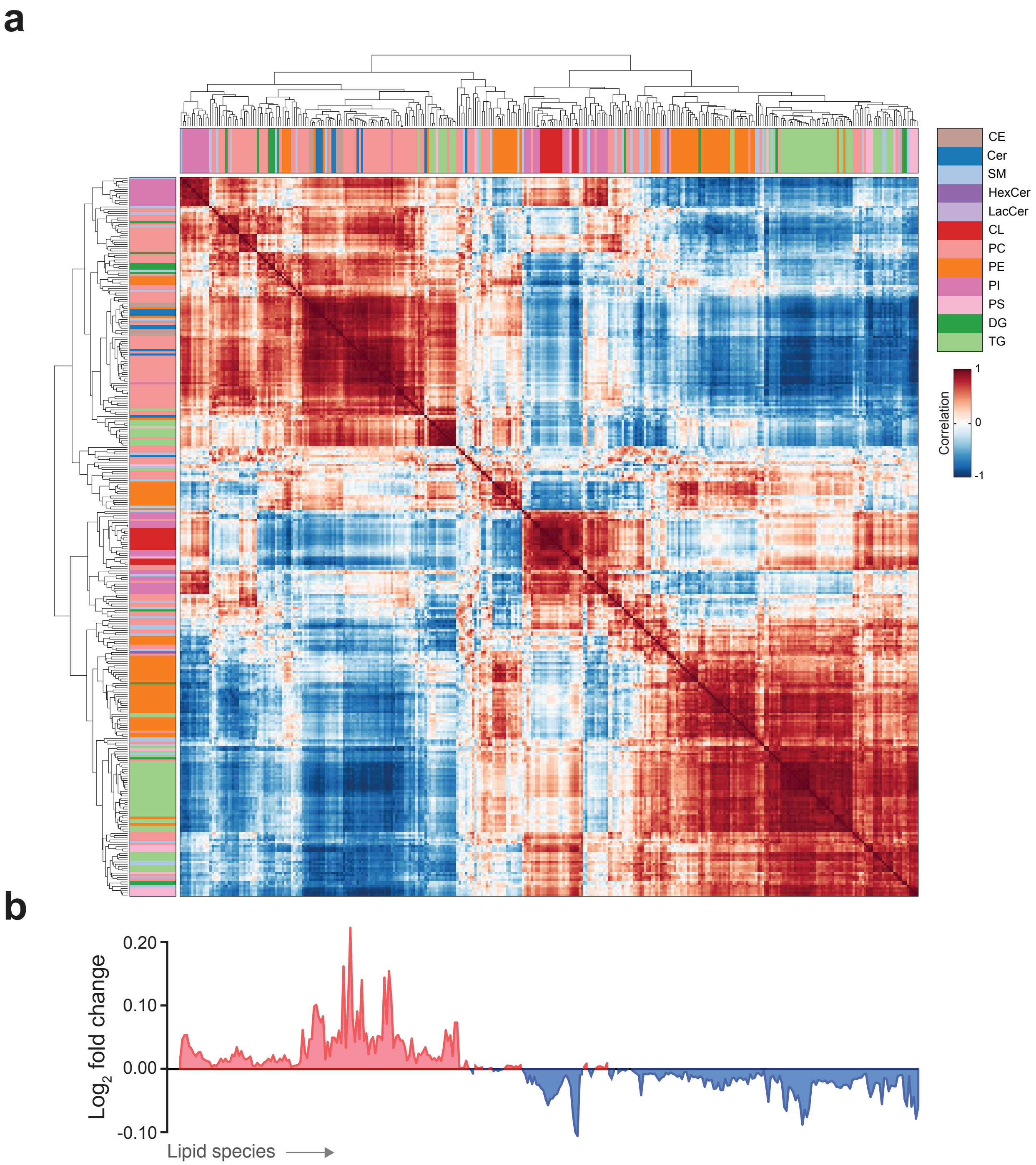


**Supplementary Fig. 2** Map of lipid correlations during ZIKV infection. **a** Correlation matrix of the 340 lipid species at 48 hpi. Each cell represents the correlation of two lipid species across the five mock and five infected samples collected 48 hpi, with direction and strength of correlation indicated by heatmap. Colored barcodes identify the subclasses of lipid species, arranged by hierarchical clustering. **b** $Log_2$ fold change in abundance of lipid species at 48 hpi, corresponding to the matrix above shown in (a). See also Supplementary Data 1.

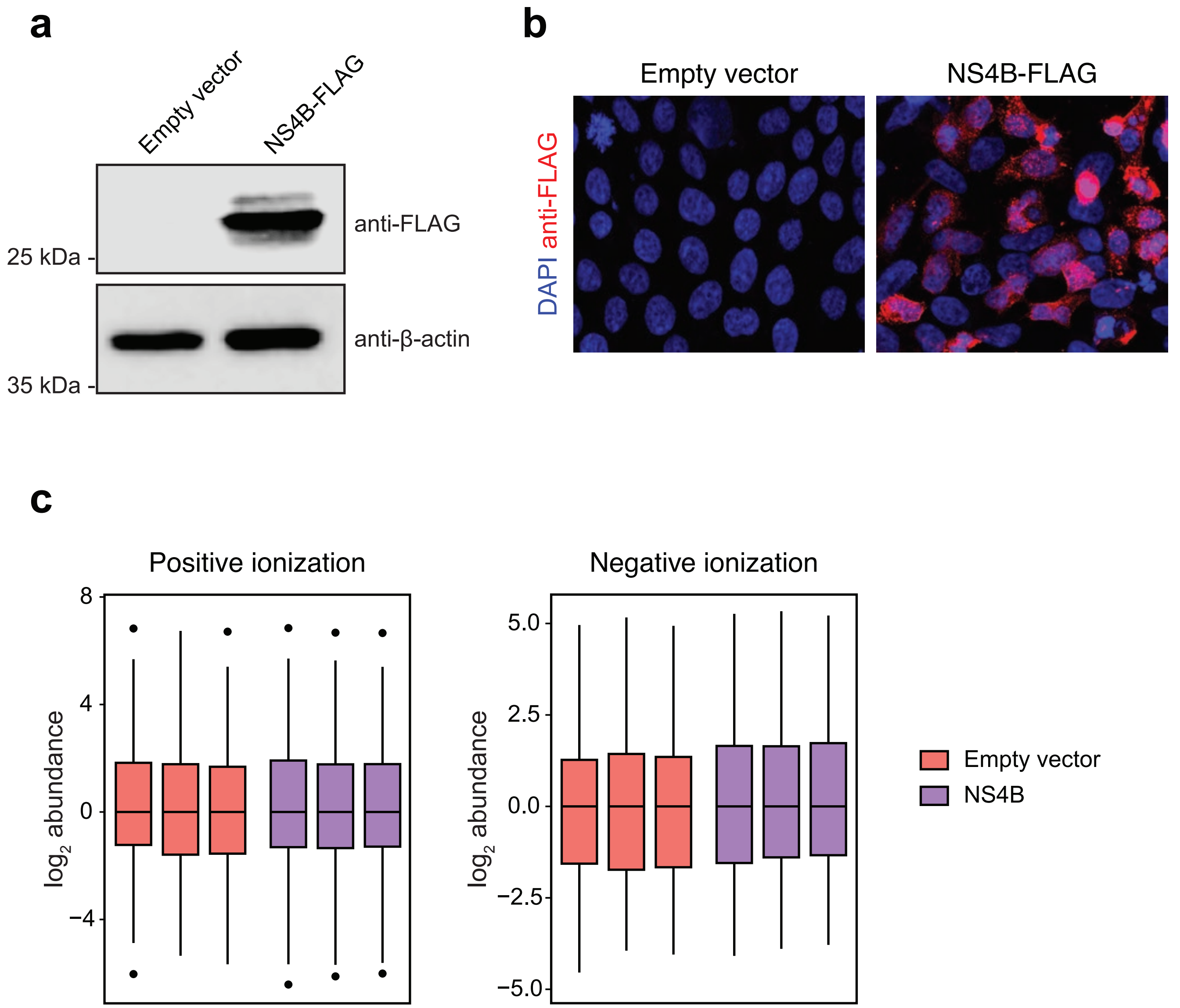


**Supplementary Fig. 3** Confirmation of NS4B expression in HEK 293T cells and lipidomics quality control. **a**, **b** HEK 293T cells were transfected with empty vector or NS4B-FLAG expression vector. NS4B expression was examined with an anti-FLAG antibody by immunoblot (a) or confocal microscopy (b). **c** Standard boxplot of the full mass spectrometry dataset after normalization. No outliers were identified. See also Supplementary Data 2.

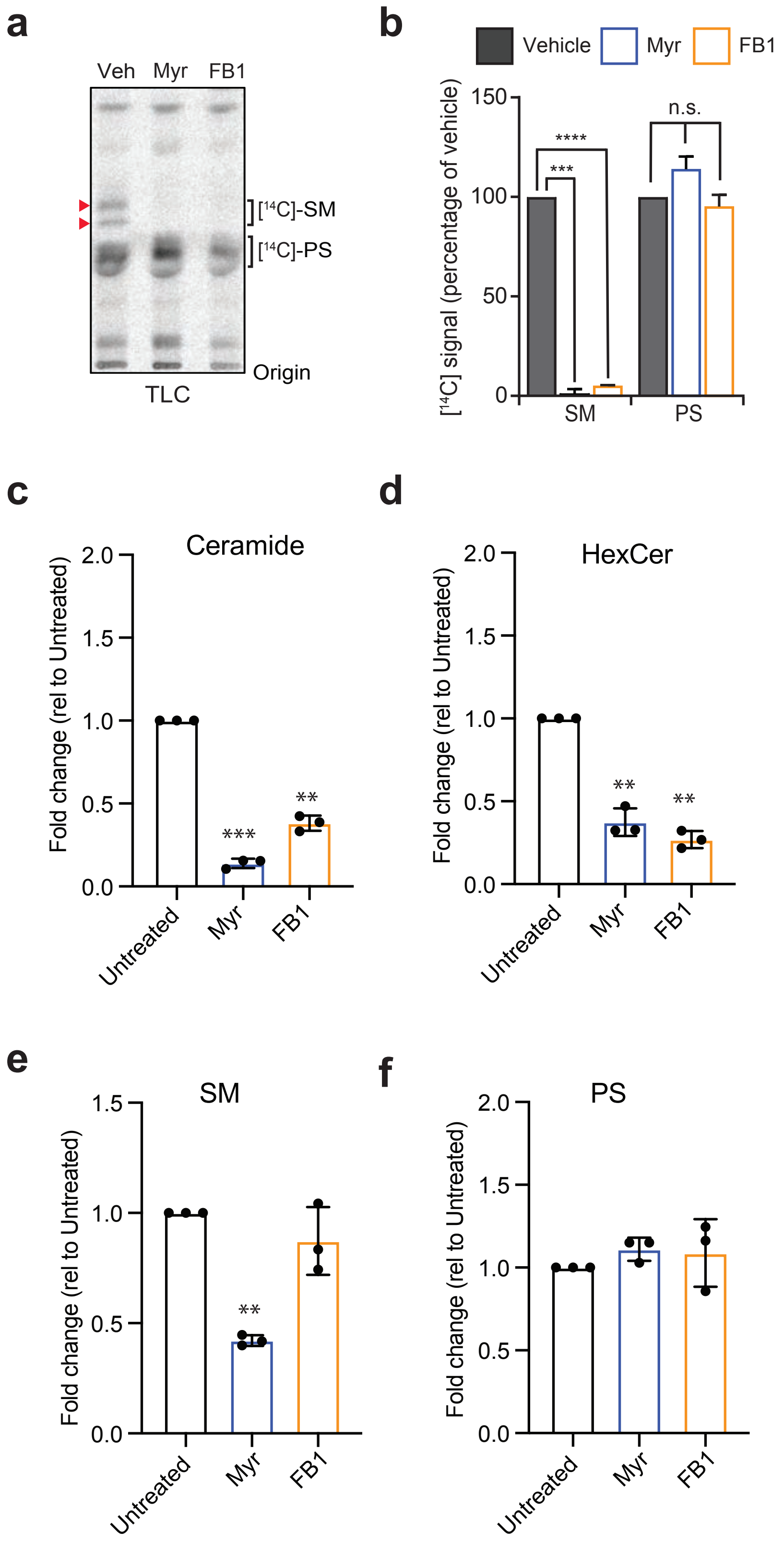


**Supplementary Fig. 4** Treatment of Huh7 cells with myriocin or FB1 specifically blocks sphingolipid biosynthesis and depletes cellular sphingolipids. **a** Huh7 cells pretreated for three days with myriocin, FB1, or DMSO (vehicle) were labelled with the sphingolipid precursor 3-L-[$^{14}$C]-serine. Total cellular lipids were extracted and resolved by TLC and $^{14}$C was visualized with autoradiography. TLC plate is representative of three independent experiments. Red arrows indicate SM bands. **b** [$^{14}$C]-SM and [$^{14}$C]-PS signals from (a) were quantified as percentage of the loading control. **c**-**f** Total lipids were extracted from control and inhibitor-treated cells and lipid profiling was performed using LC/MS. Levels of ceramide (**c**), Hexosylceramide (HexCer; **d**) sphingomyelin (SM; **e**) and phosphatidylserine (PS; **f**) are shown; $n$ = 3 biological replicates). **$P$ < 0.01, *** $P$ < 0.001, two-tailed Student's $t$-test.

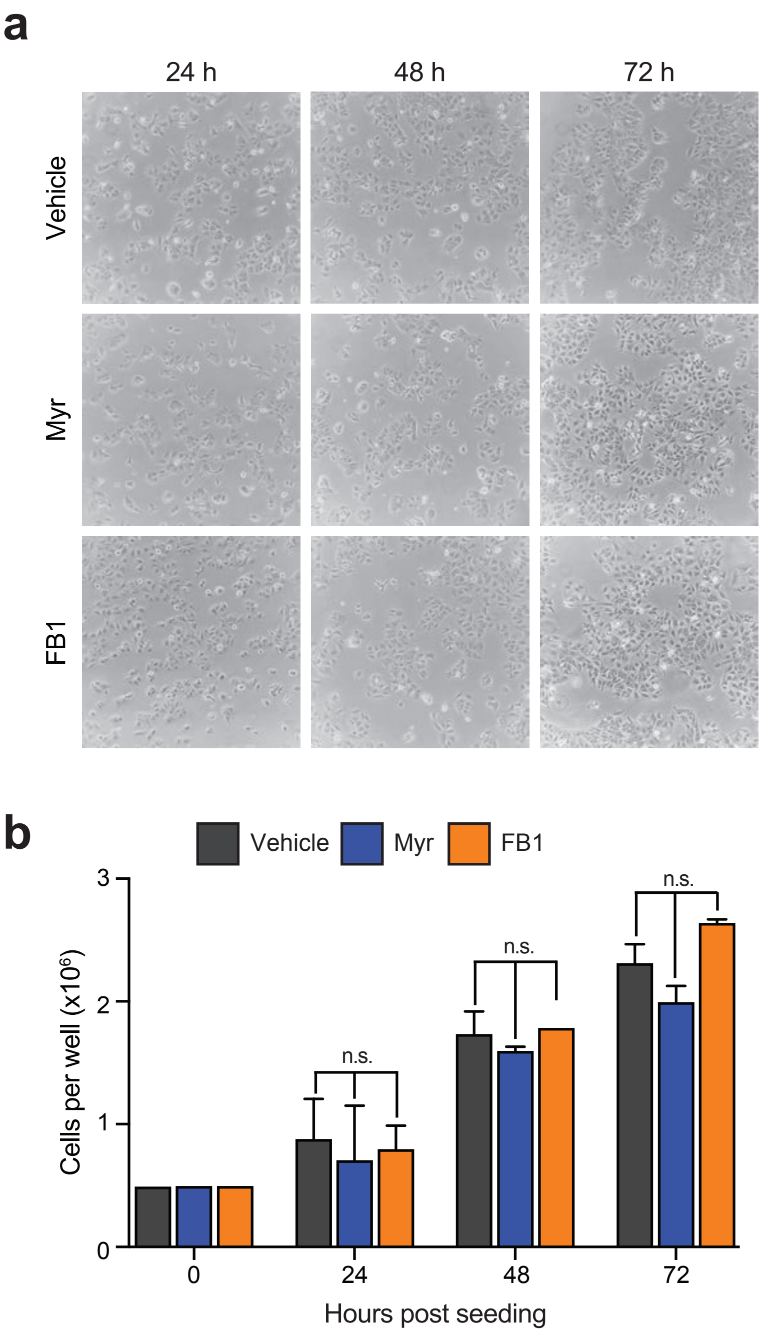


**Supplementary Fig. 5** Treatment of Huh7 cells with myriocin or FB1 does not affect cell growth or morphology. **a** Representative images of Huh7 cells treated for the indicated times with 30 μM myriocin, 5 μM FB1, or a vehicle control. **b** At 24, 48, and 72 hrs after seeding at a density of 50,000 cells/well in a 6-well plate, cells were trypsinized and counted with a hemocytometer. Data are mean ± SD. *n* = 2 independent experiments, each performed in triplicate. n.s., not significant, unpaired two-tail t test. See also the Source Data file.

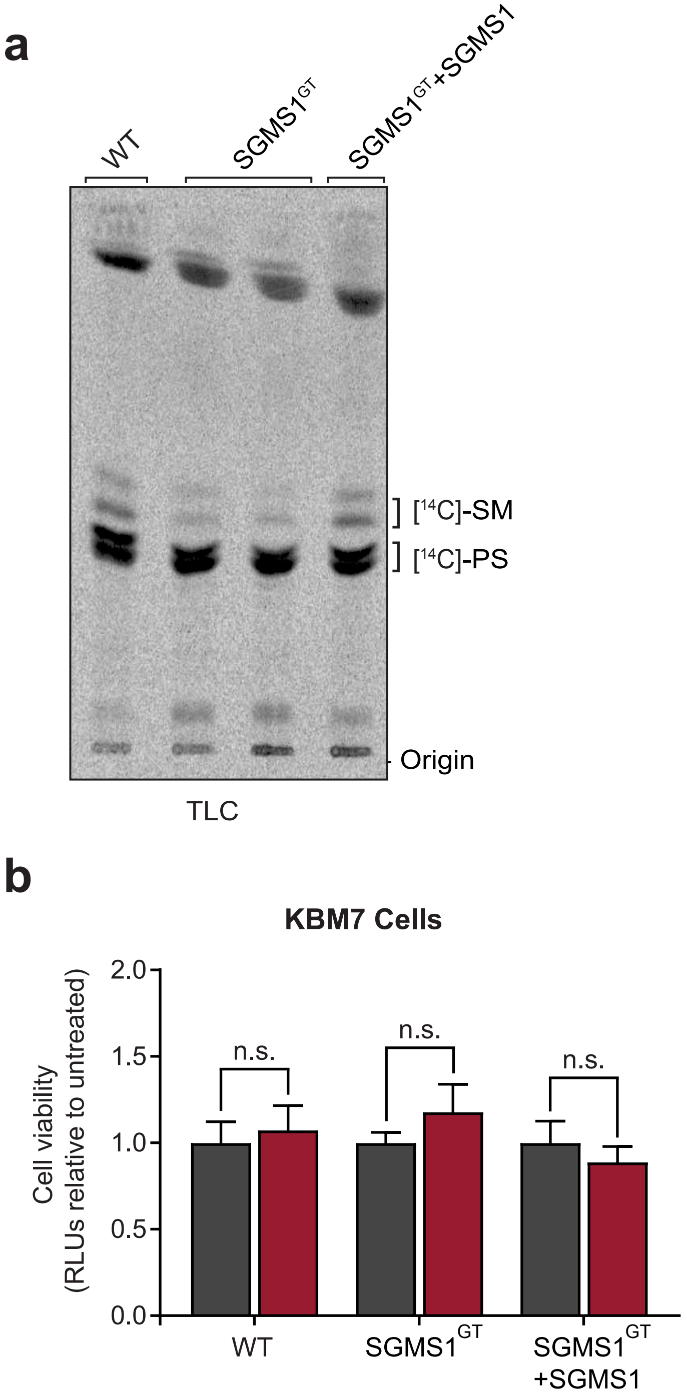


**Supplementary Fig. 6** SGMS1$^{GT}$ KBM7 cells produce reduced levels of SM, and treatment of these cells with GW4869 does not affect cell viability. **a** Wild type, SGMS1$^{GT}$ and SGMS1$^{GT}$+SGMS1 cells were labelled with the sphingolipid precursor 3-L-[$^{14}$C]-serine. Total cellular lipids were extracted and analysed by TLC and autoradiography. TLC is representative of three independent experiments. **b** The three KBM7 cell lines were treated with 10 μM GW4869 for 24 hrs, then tested for viability compared to untreated cells by measuring ATP content ($n$ = 9 replicate wells from two independent experiments). Data are mean ± SD; n.s., not significant, two-tailed Student's $t$-test.

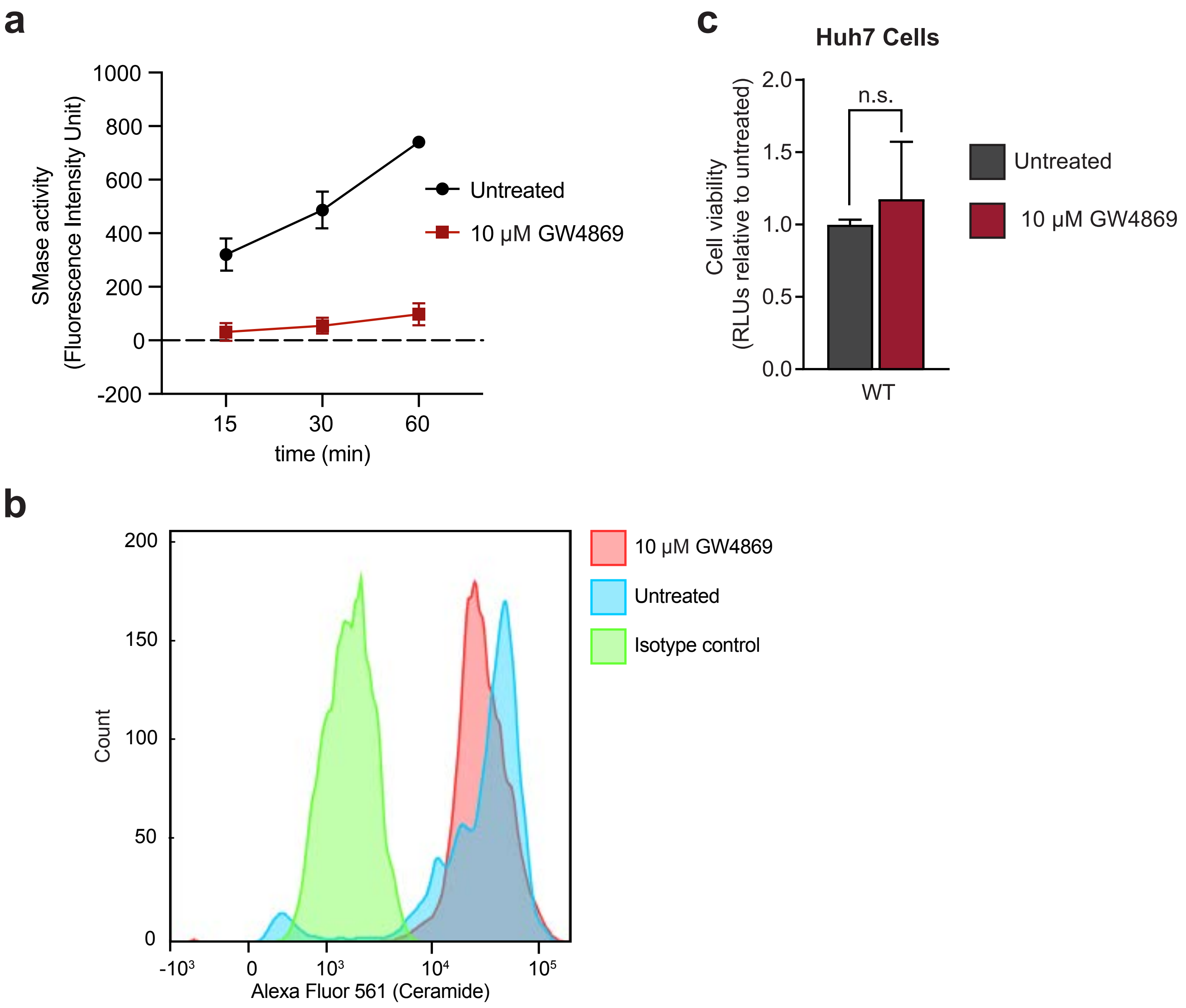


**Supplementary Fig. 7** GW4869 inhibits cellular SMase activity. **a** Huh7 cells were treated with 10μM GW4869 for 24 hrs, and the activity of cellular SMase was assayed using an Amplex Red sphingomyelinase kit (see Methods). **b** Huh7 cells were treated with GW4869 as in (a), stained with anti-ceramide monoclonal antibody, and analyzed by flow cytometry. c Huh7 cells were treated with 10 μM GW4869 as in (a), then tested for viability with the CellTiter Glo kit (n = 9 replicate wells from two independent experiments). Data are mean ± SD; n.s., not significant, unpaired two-tail t test. See also the Reporting Summary and Source Data file.

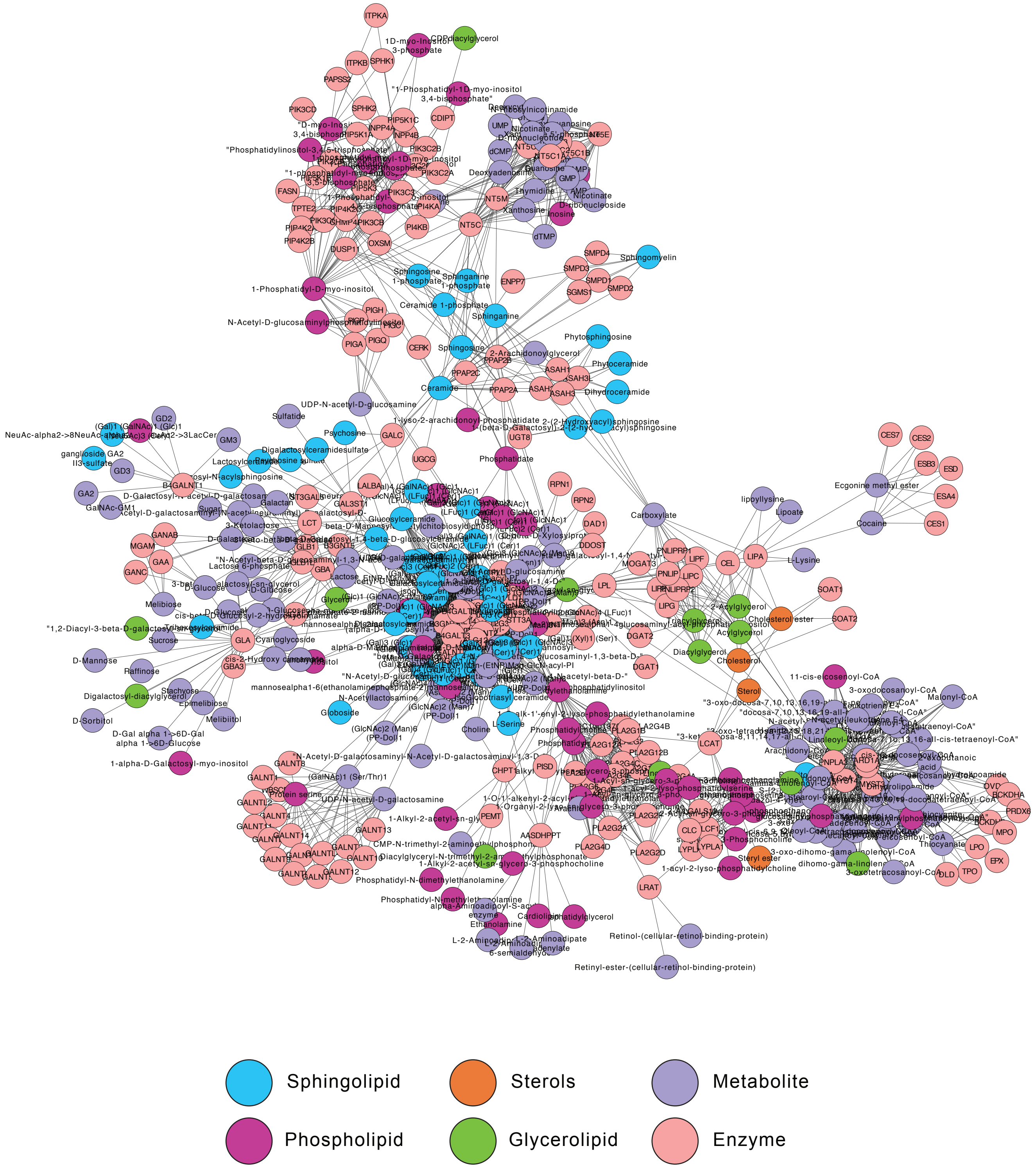


**Supplementary Fig. 8** Map of lipid metabolic network. A model of the metabolic network producing the lipid subclasses identified through lipidomics was generated in Cytoscape using the MetScape 3 plugin. Lists of genes and compounds making up the network for each lipid subclass were accessed in MetScape 3 from the KEGG database.
